# Plant leaf tooth feature extraction

**DOI:** 10.1101/418293

**Authors:** Hu Wang, Di Tian, Chu Li, Yan Tian, Haoyu Zhou

**Affiliations:** China Shipbuilding Industry Corporation, No.722 Research Institute, Wuhan 430205, Peoples R China.; Wenhua College, Faculty of Information Science and Technology, Wuhan 430074, Peoples R China.; Huazhong Univ Sci & Technol, Elect & Informat Engn Dept, Wuhan 430074, Peoples R China.

**Keywords:** leaf tooth features, leaf structure system, digital image processing

## Abstract

Leaf tooth can indicate several systematically informative features and is extremely useful for circumscribing fossil leaf taxa. Moreover, it can help discriminate species or even higher taxa accurately. Previous studies extract features that are not strictly defined in botany; therefore, a uniform standard to compare the accuracies of various feature extraction methods cannot be used. For efficient and automatic retrieval of plant leaves from a leaf database, in this study, we propose an image-based description and measurement of leaf teeth by referring to the leaf structure classification system in botany. First, image preprocessing is carried out to obtain a binary map of plant leaves. Then, corner detection based on the curvature scale-space (CSS) algorithm is used to extract the inflection point from the edges; next, the leaf tooth apex is extracted by screening the convex points; then, according to the definition of the leaf structure, the characteristics of the leaf teeth are described and measured in terms of number of orders of teeth, tooth spacing, number of teeth, sinus shape, and tooth shape. In this manner, data extracted from the algorithm can not only be used to classify plants, but also provide scientific and standardized data to understand the history of plant evolution. Finally, to verify the effectiveness of the extraction method, we use leaf tooth features and simple linear discriminant analysis to classify leaves; the results show that the proposed method achieves high accuracy as compared to other methods.

## Introduction

The potential value of leaf structure research in paleobotany, ecology, paleoecology, plant systematics, and conservation biology has begun to attract attention [1]. Generally, leaf structure allows closely related taxa to be distinguished from one another [2][3][4]. Moreover, it is found that the leaf architectural characteristics within the framework of molecular phylogenetic analysis can further highlight some evolutionary trends across angiosperms [5]. According to the biological characteristics of plant leaves, leaf structure has been examined and refined, and a relatively complete leaf structure system has been formed. In 2012, the Manual of Leaf Architecture provided a clearly defined and legendary terminology system for related research that could support wider use of leaf structure characters [6]. As one of the characteristics of plant leaves, leaf tooth is an important basis for the classification of plant leaf structure. A leaf tooth contains many characteristics of plant systematics and are extremely useful for circumscribing fossil leaf taxa [5] [7][8]. Their prevalence in fossil floras provides reliable proxy data about pre-Quaternary terrestrial paleotemperatures [9][10][11]. The size and shape of the leaf teeth are effective parameters for determining the accuracy of paleoecology analysis and paleoclimatic inference from fossil floras [12][13]. Early studies have shown that species in colder regions have more leaf teeth, while plant species in warmer regions have smoother leaf edges and fewer leaf teeth [14][15]. Therefore, fossil remains of leaves can be used as an “ancient thermometer” to help simulate past climate [10][16].

Research on leaf features is basically divided into two categories: leaf shape and leaf margin (leaf edges). In a study on leaf shape, Xiaofeng et al. used Hu geometric moment and Zernike vertical distance to measure the shape of an input leaf image for automatic leaf classification [17]; Du et al. extracted eight characteristics such as the aspect ratio, rectangular and convex area of the leaf, convex perimeter ratio, sphericity, roundness, eccentricity, and shape factor [18]. In a study on leaf edge, Zheng et al. extracted the leaf edge by using the Harris and SUSAN algorithm and calculated three characteristic parameters: leaf edge sawtooth number, sharpness, and skewness [19]; Corney automatically acquired the shape and size of the leaf tooth by identifying the tooth on the leaf edge [20]; Jin proposed a method for classifying plant leaves using the sparse matrix of leaf tooth features [21]; Chen used the ratio between the internal distance of the leaf and the Euclidean distance to represent the local concavity and convexity of leaves, and classified the whole edge, tooth edge, wave edge, and leaf crack edge; Li used data dimensionality and character weighted semi-supervised clustering algorithm to identify the leaf by synthesizing leaf shape, leaf edge, texture, and other characteristics, which could be used for several applications [23].

It should be noted that most of the extracted features are not strictly defined in botany, resulting in the inability to adopt a uniform standard to compare the accuracies of various feature extraction methods. In this study, the leaf tooth characteristics of plants are described and measured as per the botanical definition in the Manual of Leaf Architecture to provide reliable and effective features for automatic classification of plants. The basic process of the algorithm in this study is as follows: first, the leaf image is preprocessed to perform smoothing and denoising to obtain a better target image. Second, the curvature scale-space (CSS) [24] algorithm is used to extract the teeth, and according to the definition in the leaf structure manual, the features of number of orders of teeth, tooth spacing, number of teeth, sinus shape, and tooth shape are extracted one by one. The data obtained through the algorithm can not only be used to classify plants, but also provide scientific and standardized data to determine the history of plant evolution.

## Preprocessing

### Image graying

The extraction of the leaf tooth does not depend on the color of the leaf; therefore, the acquired color image is first converted to a grayscale image. The color image has three channels, R, G, and B, while the grayscale image has only one channel. After the image is grayed out, the overall and partial chromaticity and brightness of the image can be better maintained, and the amount of calculation can be drastically reduced. In this study, we use the weighted average method to convert the image to grayscale [25]. The weighted sum of the three channels of color images is expressed as follows:

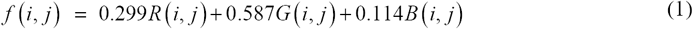

### Image segmentation

The purpose of image segmentation is to separate the leaves from the background to form a binary image for subsequent calculation of shape and texture characters. Here, the maximum interclass variance method (abbreviated as OTSU) is used to binarize the image [26]. The principle of this method is mainly to distinguish the background and target in the image by the grayscale characteristics of the image, and to calculate the interclass variance of the background and the target. The larger the variance, the greater the discrimination between the background and the target. Therefore, the segmentation with the largest variance between classes has the least probability of resulting in a misclassification.

### Image smoothing

After the image is binarized, there may be voids and breaks in the image, which may hinder the integrity of leaf extraction. Therefore, expansion, corrosion, opening, and closing operations are performed to fill small holes and gaps in the image, and finally the image is smoothed and denoised. Expansion and corrosion are the two most important and simple operators in mathematical morphology. Morphological expansion can connect the edges of a broken image to represent the complete contour of the target, while morphological erosion can remove the interference noise in the image, thereby highlighting the target.

## Leaf tooth extraction algorithm

Unlike other parts of the leaf, the characteristics of the leaf teeth are reflected in the particularity of the morphological structure, that is, leaf teeth have obvious convex and concave points, which appear as corner points on the image. Therefore, a general grading and stepwise refinement method to extract the leaf teeth can be adopted. First, the corner points (both convex and concave points in the leaf teeth) are extracted from the leaf image, and then the two are further identified according to the different characteristics of convex and concave points. The basic flowchart is shown in Fig. 1.

**Fig. 1.**
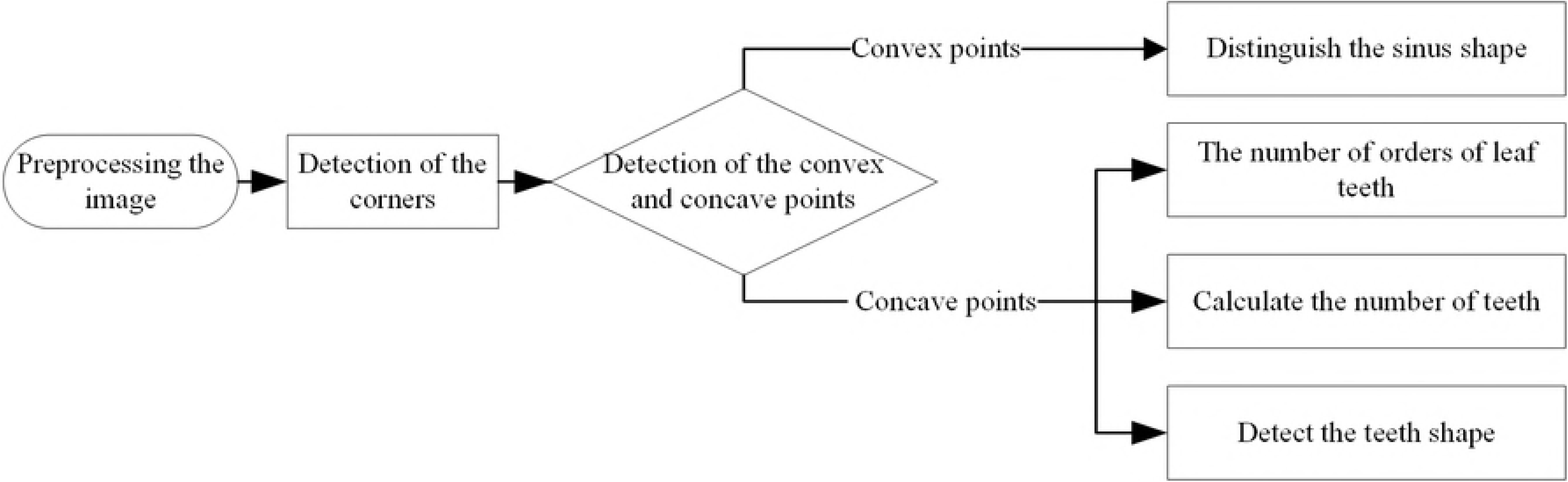
Flowchart of the leaf tooth feature extraction scheme.

### Corner detection algorithm based on multiscale curvature space

Multiscale curvature space corner detection is performed as follows [27]: (1) Applying the Canny operator to the pre-processed image to extract the leaf edge; (2) Representing the leaf edges by a single-pixel curve and marking the T-shaped corner point; (3) Calculating the curvature at all points on the contour curve with the maximum value *δ*, and defining the point where the local curvature maximum is greater than threshold t as the candidate corner point (the value of t is 162); (4) Locating the extracted corner point with the minimum value *δ*; 5) Comparing the T-shaped corner points with the candidate corner points in step 3, and eliminating two corner points that are close to each other. The basic flowchart is shown in Fig. 2.

**Fig. 2.**
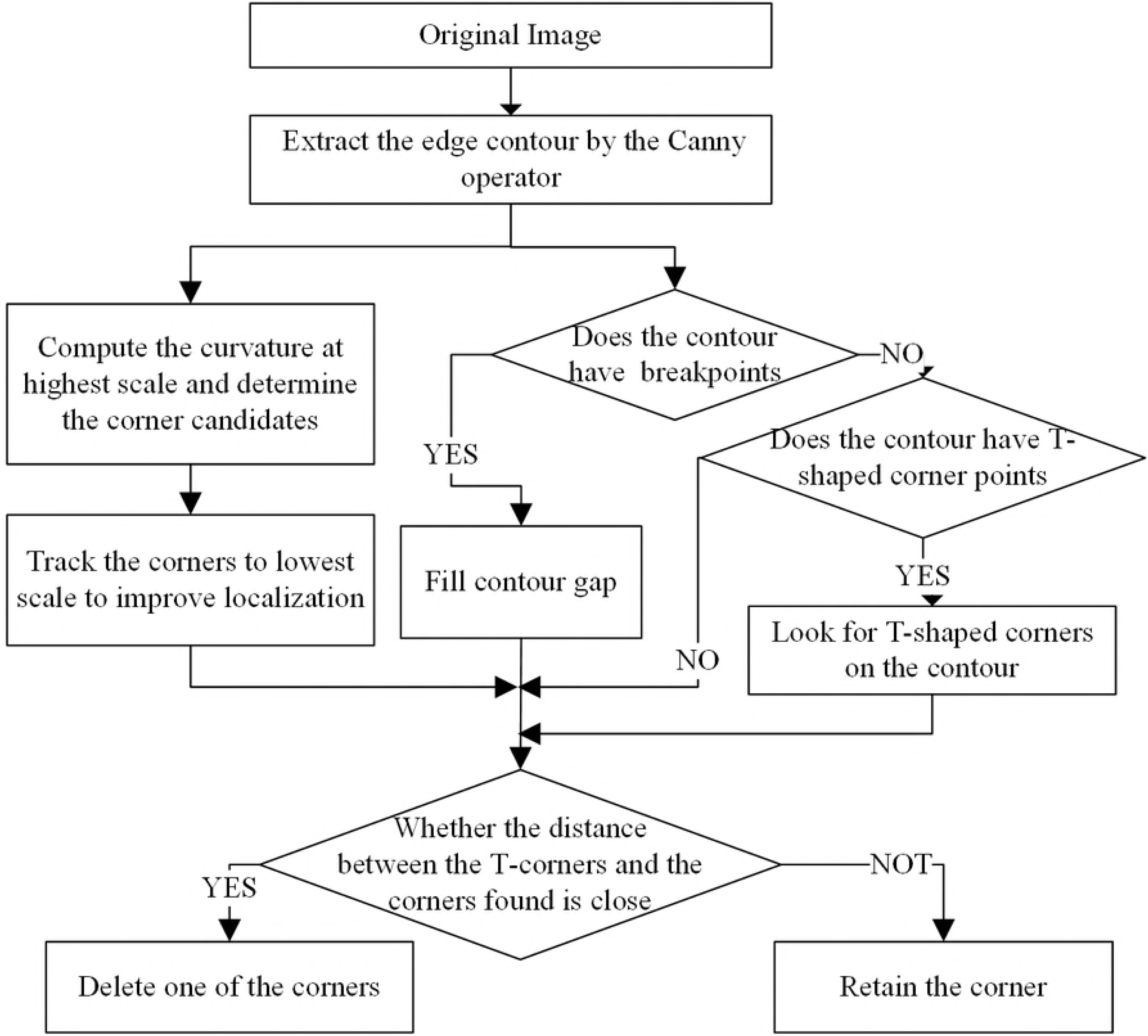
Flowchart of the CSS algorithm.

Because the value of threshold *δ* has the greatest influence on the extraction of the leaf teeth using this method, we tested the detection of the leaf teeth for 8 kinds of leaves with threshold values of 2, 4, 5, 6, and 7 (the leaf teeth of a and b are the most dense, while those of g and h are the most sparse). See Table 1 for details (other parameters are used as in [28][29] C = 1.5; T_angle = 162; sig = 4; H = 0.25; L = 0; Endpoint = 1; Gap_size = 3)

**Table 1.**
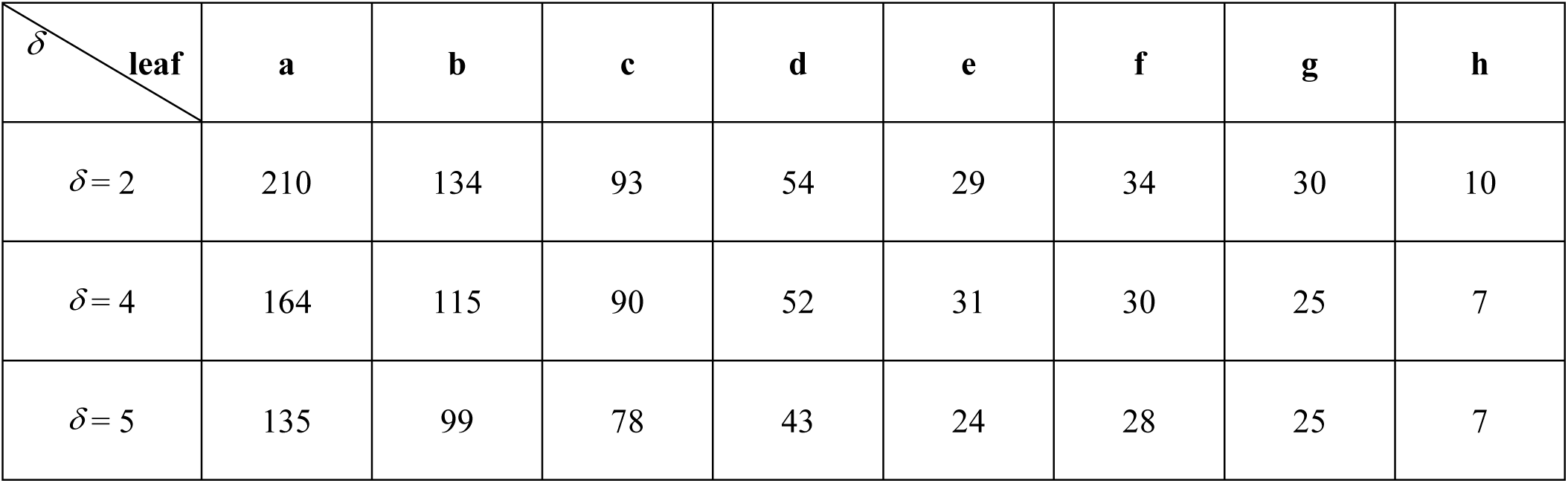
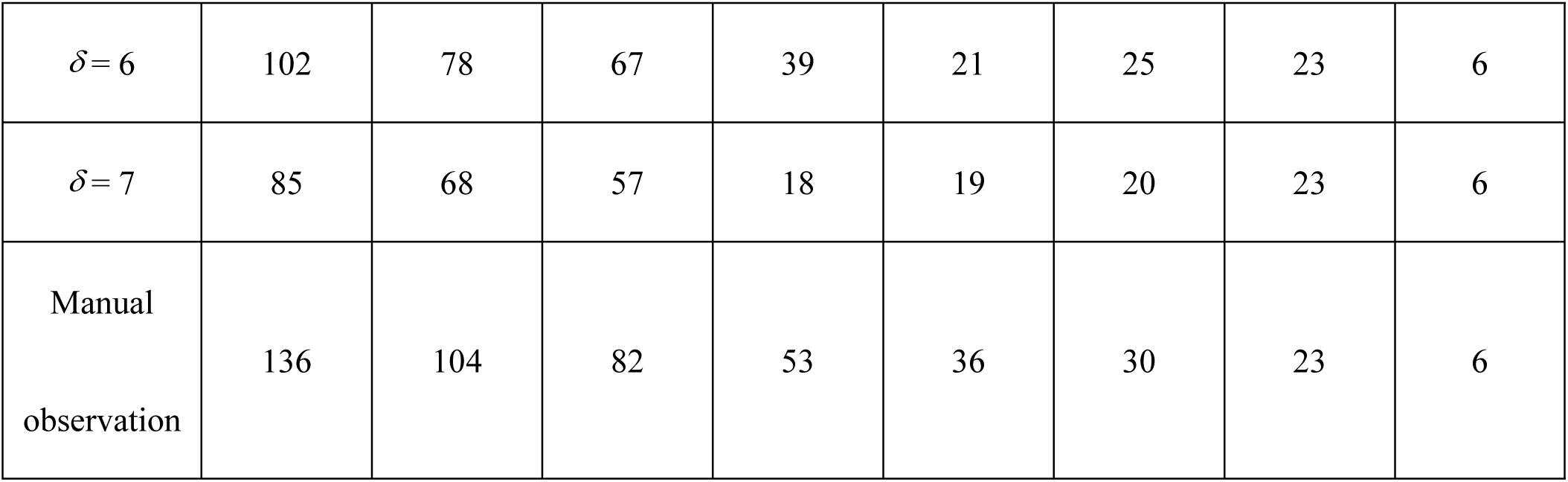
Relationship between the number of extracted leaf teeth and the value of *δ*.

It can be seen from Table 1 that the value of *δ* affects the accuracy of leaf tooth extraction. The number of leaf teeth detected by this method decreases as the value of *δ* increases. For different types of leaves, the value of *δ* is different corresponding to the smallest extraction error. For example, leaves a and b with dense teeth have the highest accuracy at *δ* =5, and leaves g and h with sparse number of teeth have highest accuracy at *δ* =6. That is, for leaves with dense teeth, the leaf tooth point is captured more accurately when the curvature scale is small, and for leaves with sparse teeth, when the curvature scale is large, a part of the noise can be shielded to obtain a more precise leaf tooth point.

### Extraction of convex leaf tooth points

The extraction of convex points can be divided into two steps: first, all the corner points are detected as the candidate points by the multiscale curvature spatial corner detection method introduced in Section 3.1. Second, considering the detected corner points may contain concave points and non-convex non-concave points, the convex points must be detected based on the different characteristics of these three types of points.

The specific detection method of the convex points is as follows:

(1) Use the CSS algorithm to detect all the corner points of the leaf;
(2) With the corner point as the center, set the radius to L, and calculate the number of target points and background points falling in the circle. If the number of target points is less than that of the background points, it is a convex point. As shown in Fig. 3(a), P_1_ is a convex point, and Fig. 3(b) is the actual image with convex points calibrated by the algorithm.

**Fig. 3.**
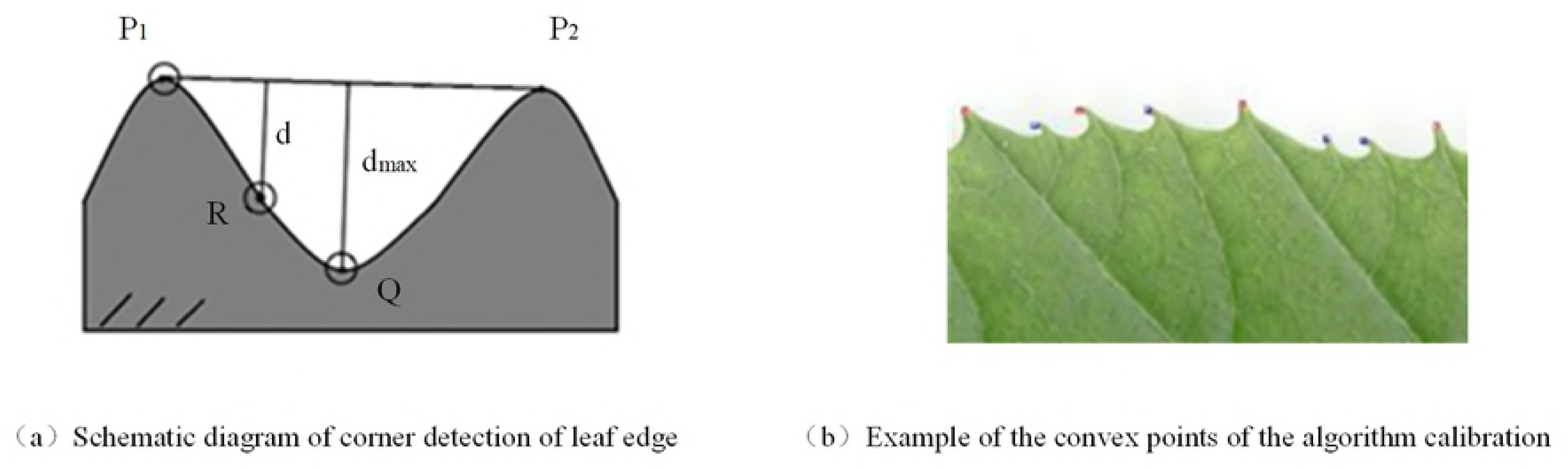
Example of convex points calibrated by the algorithm.

As shown in Fig. 3(a), the method of extracting the leaf tooth concave points is as follows: P_1_ and P_2_ are two adjacent convex points, the concave point Q will be between the two convex points, and the distance d_max_between Q and the line P_1_P_2_ is inevitably greater than the distance d between the other points on the curve between P1 and P2. Therefore, by calculating the distance d from the point on the curve to the straight line determined by the two adjacent convex points, and comparing it, point Q corresponding to the maximum distance dmax is found to be a concave point.

### Extraction of leaf tooth features

According to the leaf structure classification system, the characteristics of the leaf teeth can be described by the number of orders of teeth, tooth spacing, number of teeth, sinus shape, and tooth shape. The description and extraction methods of these features will be given one by one.

### Leaf tooth feature description

(1) The number of orders of teeth is the number of discrete sizes of teeth. Order of one means that all teeth have the same or continuous change in size, as shown in Fig. 4(a). Order of two means teeth have two distinct sizes, as shown in Fig. 4(b), and so on.

**Fig. 4.**
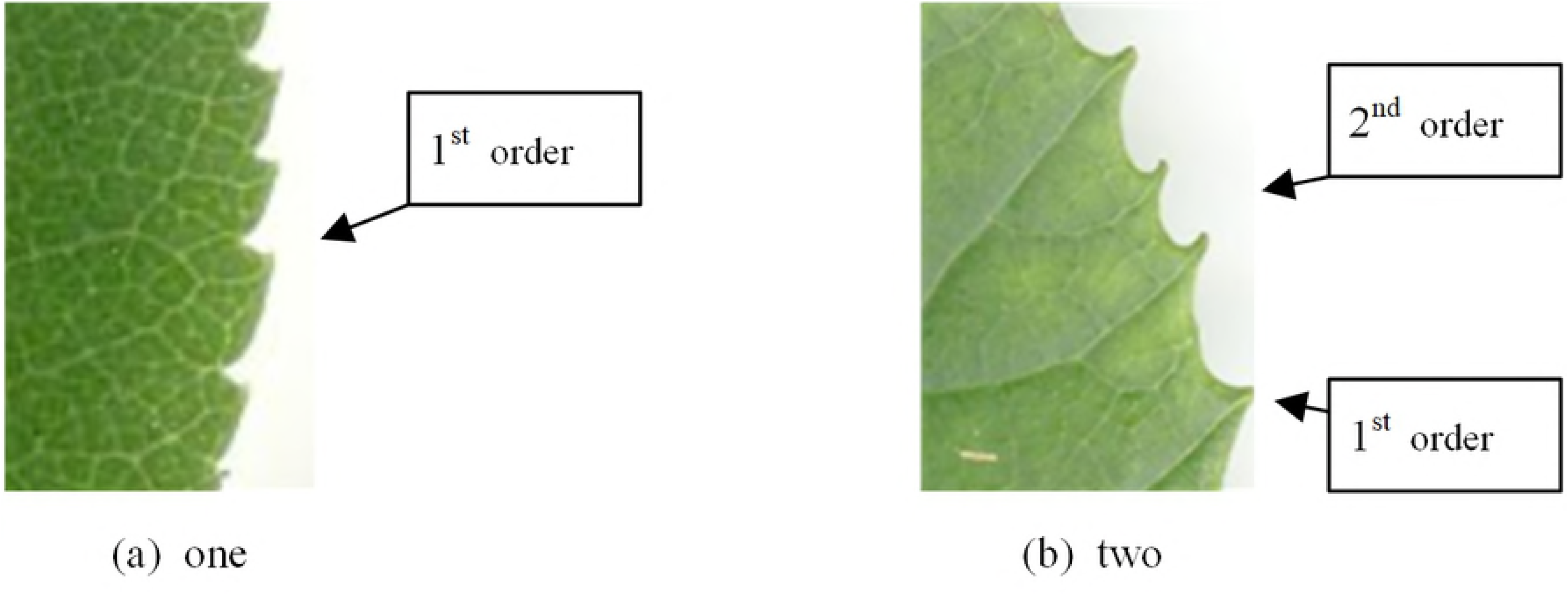
Schematic diagram of the orders of leaf tooth.

(2) The tooth spacing refers to the distance between adjacent teeth. Here, we just perform qualitative analysis, not quantitative analysis; if the minimum intertooth distance > 60% of the maximum intertooth distance, the tooth spacing is considered to be regular, otherwise it is considered to be irregular [6].

(3) The number of teeth is the total number of teeth on a leaf.

(4) Sinus shape is the shape of a concave edge on the leaf such as a horn or a rounded shape, as shown in Fig. 5.

**Fig. 5.**
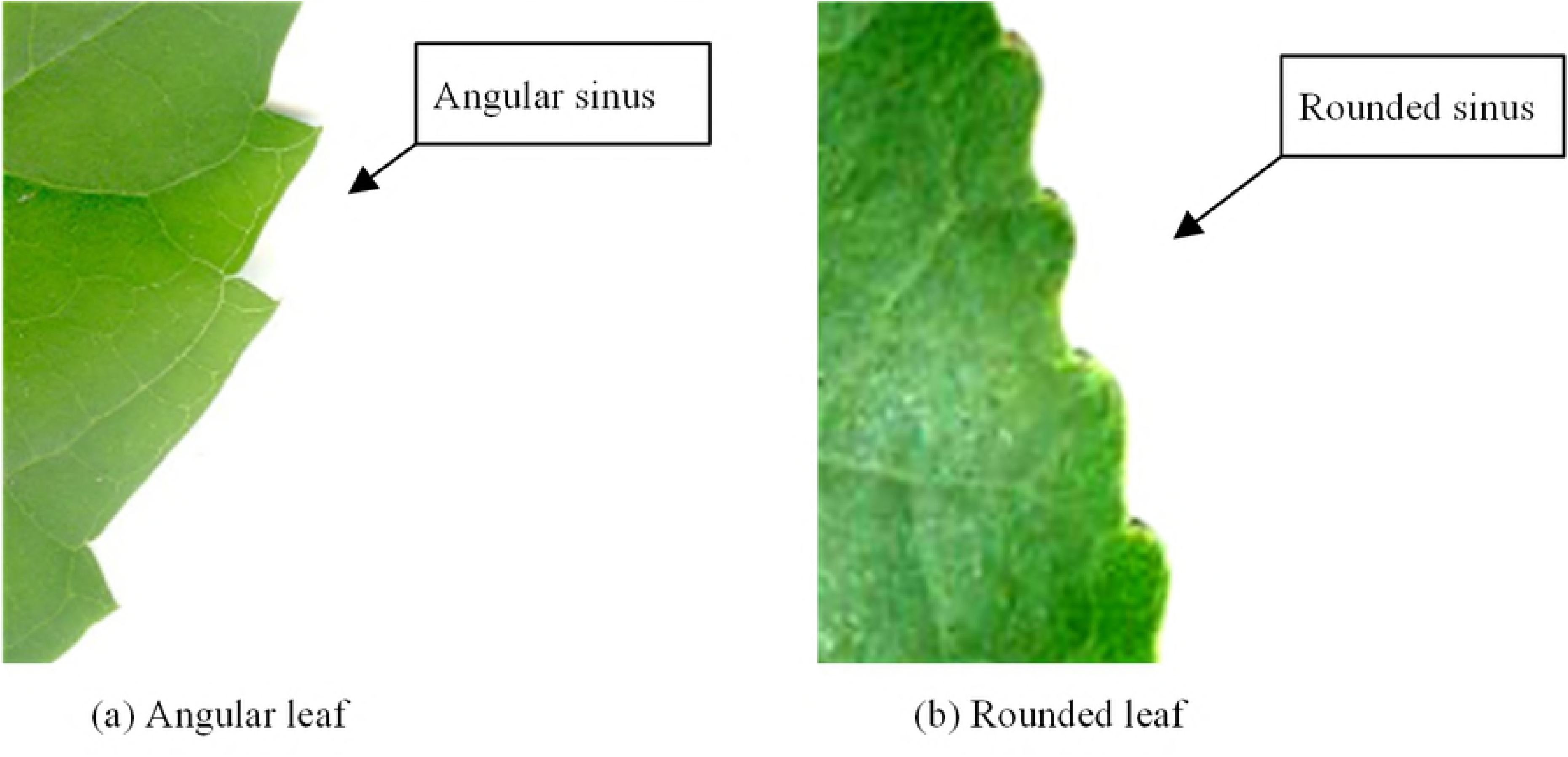
Sinus shape.

(5) The tooth shape is the distal and proximal flank curvatures relative to the midline of the tooth. The following states and abbreviations are used: convex(cv), straight(st), concave(cc), flexuous(fl), and retroflexed(rt) [6], as shown in Fig. 6.

**Fig. 6.**
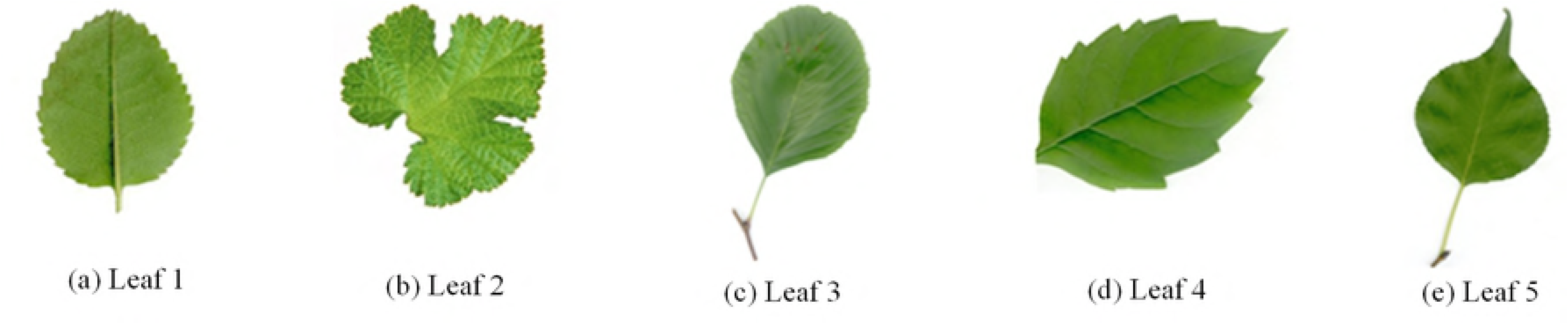
Chart of possible tooth shapes.

### Leaf tooth feature extraction

#### Number of orders of teeth

1. Calculate the distance from the convex point to the line formed by adjacent concave points to indicate the length of the teeth;

2. Normalize the teeth length, that is, divide the distance by the length of the longest tooth in the leaf (after removing abnormal points, which have values greater than twice the mean of the teeth length);

3. After normalization, the number of teeth for which the tooth length is less than the half of the maximum length are counted. If they are greater than half of the total number of leaf teeth, the leaf is considered to have two orders of teeth, otherwise it is considered to have the order of one;

4. If step 3 shows that the leaf has two orders of teeth, it is necessary to distinguish the first-order teeth and the second-order teeth by comparing the teeth length. The length of the first-order leaf teeth is significantly larger than the length of the two adjacent leaf teeth and the second-order teeth do not have this characteristic, the teeth can be distinguished.

#### Tooth spacing

1. Calculate the tooth spacing, that is, the distance between adjacent teeth;

2. Compare the minimum intertooth distance d_min_ with the maximum intertooth distance d_max_. If d_min_>(d_max_*0.6), the tooth spacing characters is regular; otherwise, it is irregular.

#### Tooth number

Calculate the total number of teeth of a leaf (i.e., the number of convex points using the abovementioned detection method).

#### Concave shape

1. Calculate the curvature at the concave point of a leaf whose concave shape is rounded;

2. Calculate the curvature at the concave point of a leaf that has an angular concave shape;

3. According to the statistical data in steps 1 and 2, determine the curvature threshold value which distinguishes the angular concave shape and the rounded concave shape.

#### Tooth shape

1. Extract the coordinates of an arc between a concave point and a convex point on the profile curve of the leaf tooth;

2. Fit the coordinates of the quadratic polynomial to obtain the corresponding polynomial;

3. Examine the quadratic parameter of the quadratic polynomial; if the parameter is greater than 0, it is convex or flexuous; if the parameter is less than 0, it is concave or retroflexed; if the parameter is equal to 0, it is straight.

## Experiments

### Experimental plan

In order to verify whether the proposed leaf structure feature description algorithm is scientific and effective, in this section, we provide relevant experimental results and analyze them.

The test data were obtained from standard plant leaf database (Swedish Leaf Dataset and Institute of Botany, Chinese Academy of Sciences) [30], and several representative shaped leaves were selected, as shown in Fig. 7.

**Fig. 7.**
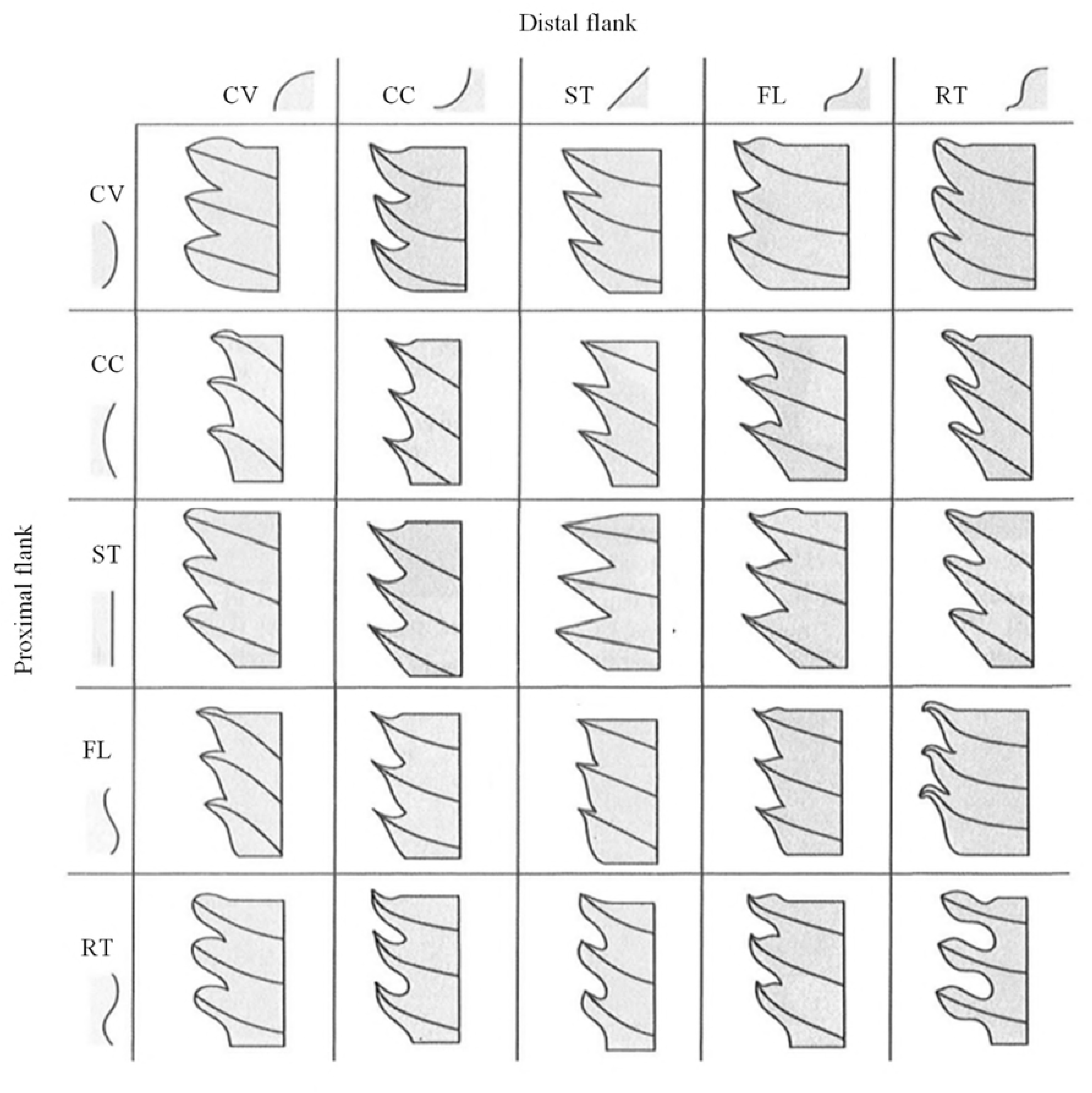
Example of test leaves.

The extraction algorithm proposed in Sections 3 and 4 in this paper was used to extract the tooth features. We verify the validity of features extracted in two ways. The first method uses a manual view (Experiment 1) to verify whether the features extracted by the method are consistent with the actual characters of the leaf; the second method applies the extracted features to a classification algorithm (Experiment 2) to indirectly verify the validity of the features by classification; the classification is performed using a general evaluation method: confusion matrix and Kappa statistics. The confusion matrix consists of four values: true positive (TP), false positive (FP), false negative (FN), and true negative (TN). Based on these data, the value of true negative rate (specificity), negative predictive value (NPV), precision, recall, accuracy, F-measure, and Kappa coefficient (Kappa) are calculated [31]. Table 2 lists the Kappa coefficients for the classification, and the accuracy criteria are determined using the method proposed by Landis and Koch [32].

**Table 2.**
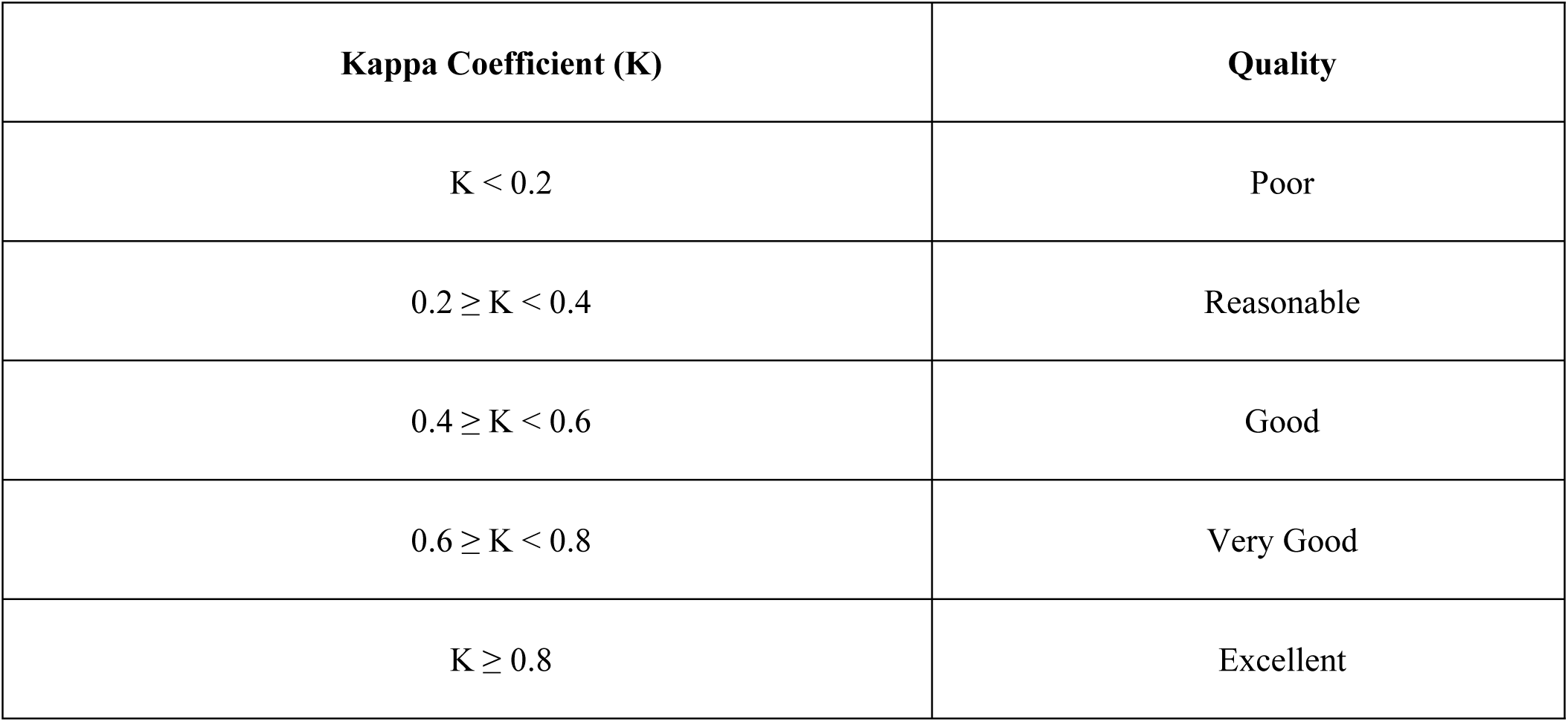
Level of classification accuracy according to the Kappa coefficient value.

| **Kappa Coefficient (K)** | **Quality** |
| --- | --- |
| $K < 0.2$ | Poor |
| $0.2 \leq K < 0.4$ | Reasonable |
| $0.4 \leq K < 0.6$ | Good |
| $0.6 \leq K < 0.8$ | Very Good |
| $K \geq 0.8$ | Excellent |

## Results and Analysis

### Experiment 1

The experimental results of the first method (manual view) of the algorithm validation are shown in Table 3-5.

**Table 3.**
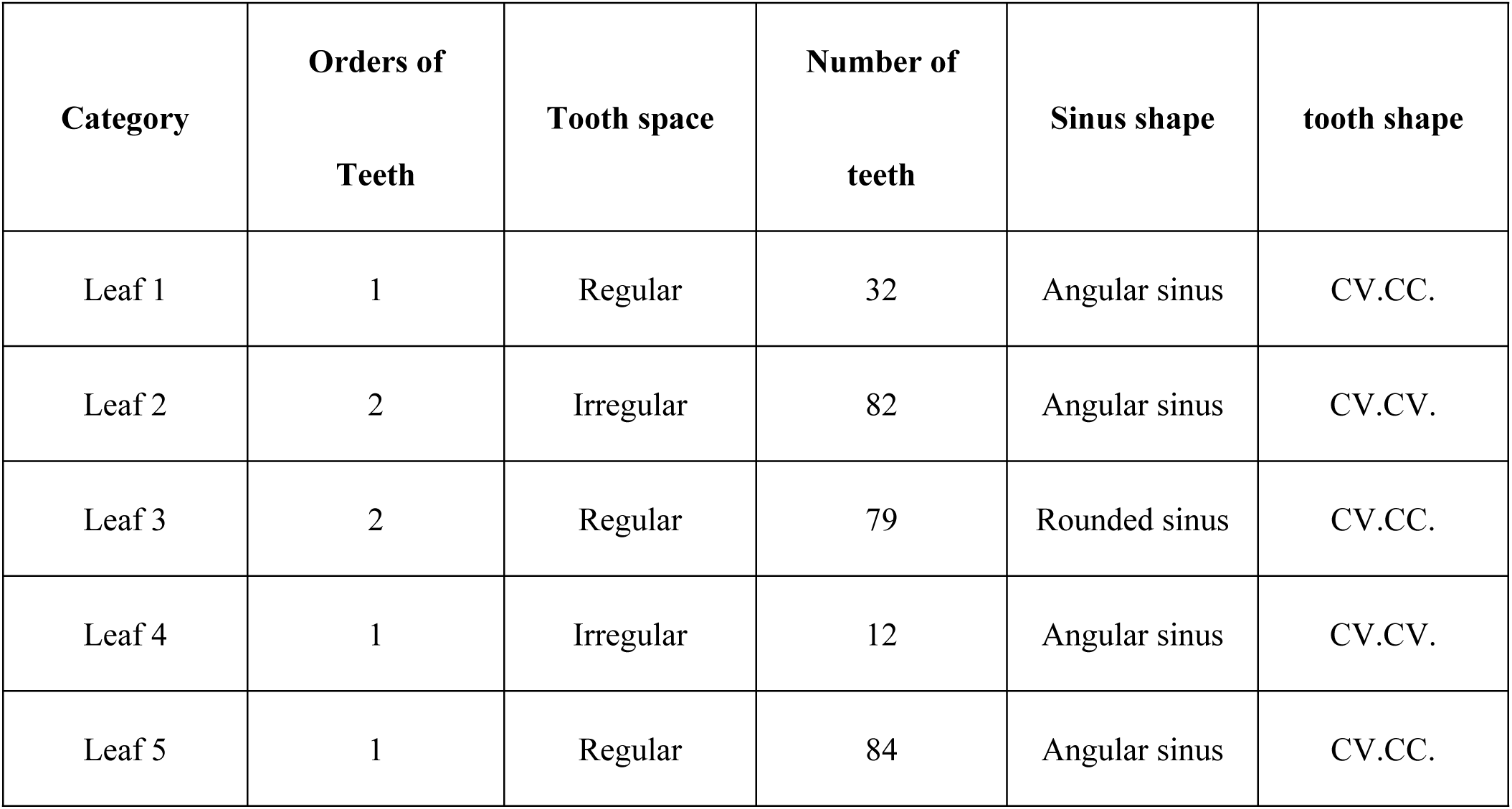
Results of tooth extraction.

**Table 4.** Accuracy statistics of the orders of leaf tooth.

| Category | First-order teeth |  | Accuracy | Second-order teeth |  | Accuracy |
| --- | --- | --- | --- | --- | --- | --- |
|  | Positive | Negative |  | Positive | Negative |  |
| Leaf 1 | 23 | 6 | 79% | - | - | - |
| Leaf 2 | 17 | 7 | 70% | 42 | 16 | 71% |
| Leaf 3 | 33 | 1 | 97% | 47 | 6 | 88% |
| Leaf 4 | 11 | 2 | 84% | - | - | - |
| Leaf 5 | 27 | 8 | 77% | 49 | 9 | 84% |

**Table 5.**
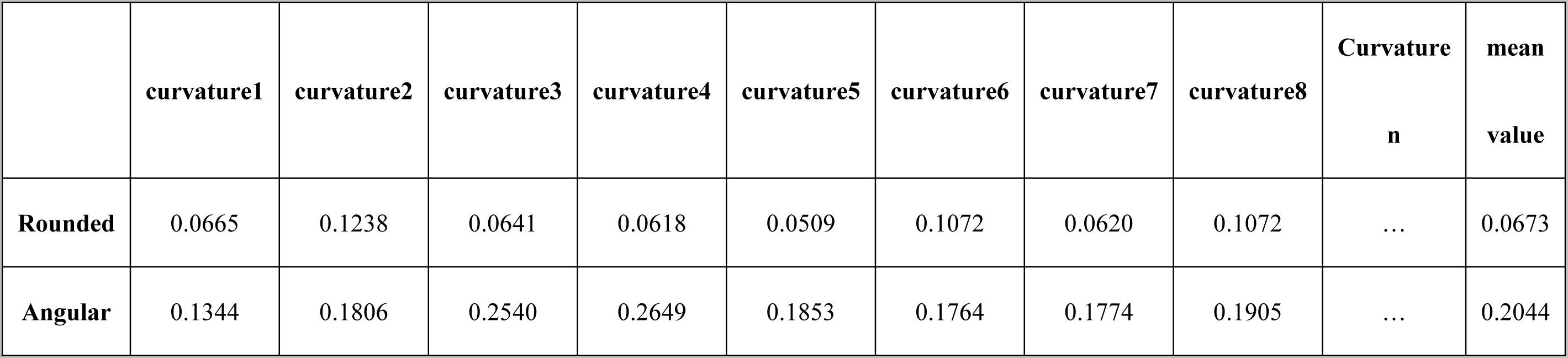
Statistics of the curvature of the concave shape.

From the experimental results listed in Table 3 – Table 5, it can be seen that the method accurately extracts the tooth characteristics of the leaf. Moreover, the tooth spacing, the number of teeth, the shape of the concave points, and the number of orders of teeth are also in line with the observed results, which indicates that the CSS method is accurate and reliable for the extraction of leaf teeth features. Simultaneous curvature is an effective method to describe the shape of leaf concave shape.

### Experiment 2

In order to further test the effectiveness of the extraction method proposed in this paper, the extracted features are used for leaf classification; thus, the effectiveness of method is verified by the accuracy of classification. Because we use the same method to extract multiple features from the same sample, if the accuracy of the classification is higher, the features have a better distinction. An extraction method of leaf tooth features has been discussed in detail and the classification of leaves based on leaf teeth characteristics has been studied, which achieved good results [20]. Other methods [21][22][23] use a combination of leaf tooth features and non-leaf tooth features; the focus of this study is only on the leaf teeth features, so the comparison method is based on the selection proposed in [20]. A leaf identification method identifies features such as total area of teeth, internal angles, number of teeth, and total length of outer edges.

The original input uses four kinds of plant leaves, a total of 700 leaf images (from the Swedish Leaf Dataset [30]; the dataset contains scanned images of 15 leaves and the selected species have greater similarity, which is considered as challenging to be classified). We use our method to extract the leaf tooth features from the original leaves as an input dataset; we also need to divide the data into a non-overlapping training set (90% of the records) used to estimate the model parameters, and a testing set (10% of the records) used to estimate its accuracy.

First, we predict the category labels using a multiclass linear discriminant analysis (LDA) classifier. The feature classification test results extracted by the proposed method and the method proposed in [20] are shown in Table 6.

**Table 6.** LDA classification results.

| Categor<br>y | Method | Specificit<br>y | NPV | Precisio<br>n | Recal<br>l | F1-measure | Accurac<br>y | Kapp<br>a |
| --- | --- | --- | --- | --- | --- | --- | --- | --- |
| 1 | Our method | 0.2667 | 0.444<br>4 | 0.3939 | 0.866<br>7 | 0.5417 | 0.4167 | 0.2222 |
|  | Proposed in<br>[20] | 0.1333 | 0.4 | 0.2667 | 0.8 | 0.4 | 0.3 | 0.0667 |
| 2 | Our method | 0.5556 | 0.446<br>4 | 0 | 0 | - | 0.4167 | 0.2222 |
|  | Proposed in<br>[20] | 0.3333 | 0.288<br>5 | 0.375 | 0.2 | 0.2609 | 0.3 | 0.0667 |
| 3 | Our method | 0.3778 | 0.34 | 0.8 | 0.533<br>3 | 0.64 | 0.4167 | 0.2222 |
|  | Proposed in<br>[20] | 0.3778 | 0.293<br>1 | 0.5 | 0.066<br>7 | 0.1176 | 0.3 | 0.0667 |
| 4 | Our method | 0.4667 | 0.446<br>8 | 0.3077 | 0.266<br>7 | 0.2857 | 0.4167 | 0.2222 |
|  | Proposed in<br>[20] | 0.3556 | 0.290<br>9 | 0.4 | 0.133<br>3 | 0.2 | 0.3 | 0.0667 |

It can be seen from Table 6 that the accuracy of leaf classification using the proposed method is 41.67%, and Kappa coefficient classified the result as “reasonable.” However, the accuracy of leaf classification is only 30% using the extraction method in [20] and the Kappa coefficient classified the result as “poor” according to the accuracy levels (K<0.2). Therefore, the feature extracted by the proposed method has higher accuracy for leaf classification; that is, the features extracted using this method have better discrimination capability for different leaves.

Second, we use a multiclass support vector machine (SVM) classifier to predict the category of leaves. The results of the feature classification test for the proposed method and the method proposed in [20] are shown in Table 7.

**Table 7.**
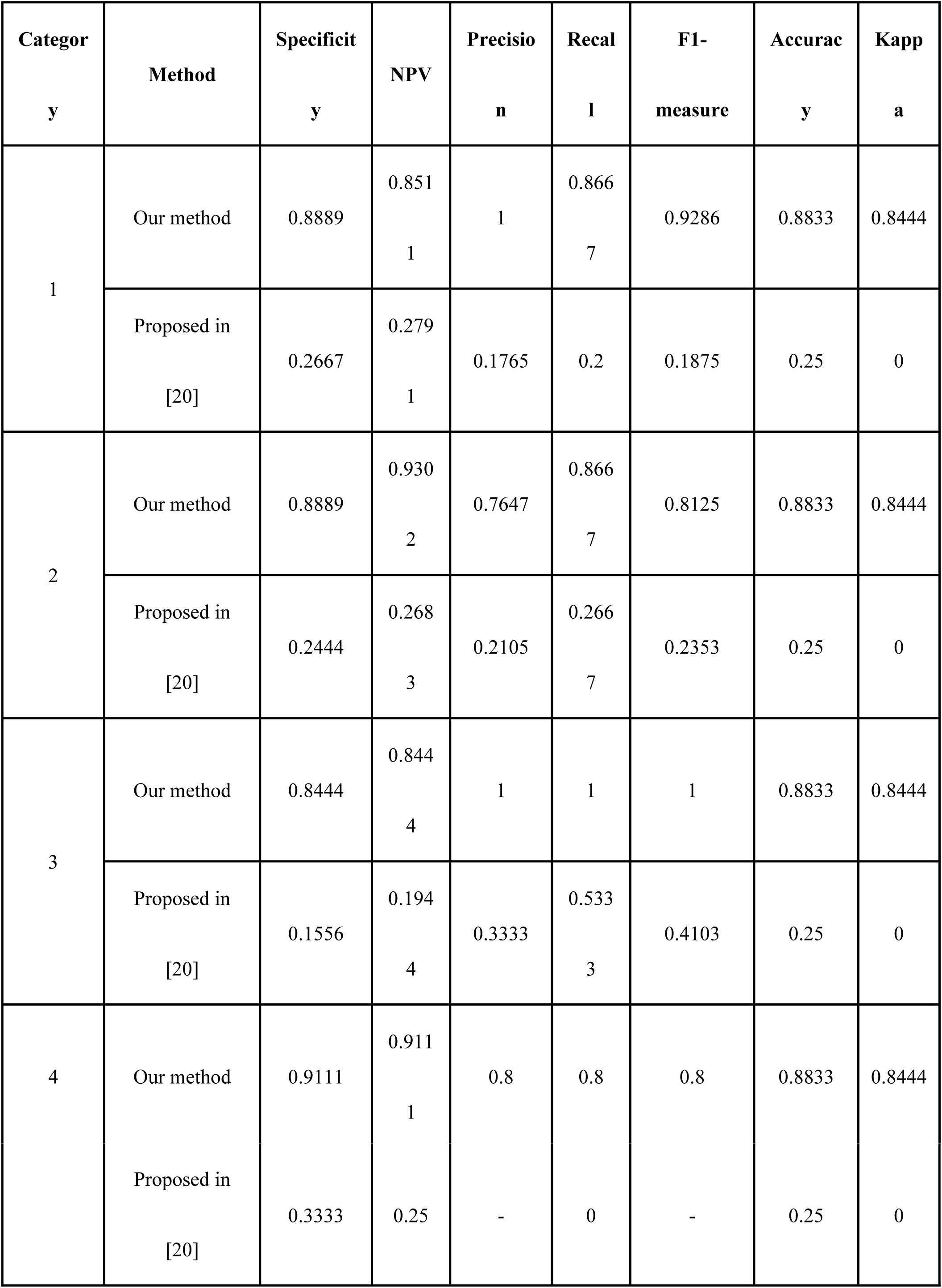
SVM classification results.

| Category | Method | Specificity | NPV | Precision | Recall | F1-measure | Accuracy | Kappa |
| --- | --- | --- | --- | --- | --- | --- | --- | --- |
| 1 | Our method | 0.8889 | 0.8511 | 1 | 0.8667 | 0.9286 | 0.8833 | 0.8444 |
|  | Proposed in [20] | 0.2667 | 0.2791 | 0.1765 | 0.2 | 0.1875 | 0.25 | 0 |
| 2 | Our method | 0.8889 | 0.9302 | 0.7647 | 0.8667 | 0.8125 | 0.8833 | 0.8444 |
|  | Proposed in [20] | 0.2444 | 0.2683 | 0.2105 | 0.2667 | 0.2353 | 0.25 | 0 |
| 3 | Our method | 0.8444 | 0.8444 | 1 | 1 | 1 | 0.8833 | 0.8444 |
|  | Proposed in [20] | 0.1556 | 0.1944 | 0.3333 | 0.5333 | 0.4103 | 0.25 | 0 |
| 4 | Our method | 0.9111 | 0.9111 | 0.8 | 0.8 | 0.8 | 0.8833 | 0.8444 |
|  | Proposed in<br>[20] | 0.3333 | 0.25 | - | 0 | - | 0.25 | 0 |

It can be seen from Table 7 that the accuracy of leaf classification with the proposed extraction method is 88.33%, and the Kappa coefficient classified the solution as “excellent”; the accuracy of leaf classification is only 25% if we use the extraction method proposed in [20], and the 4th leaf is classified incorrectly, for which the Kappa coefficient classified the results as “poor.” This proves that the leaf tooth features as referred to leaf structure classification system in botany, such as number of orders of teeth, tooth spacing, tooth number, concave shape and so on, can accurately describe the differences between different types of plant leaves; the accuracy of SVM classification using this method is significantly higher than that using LDA method.

## Discussion

The extraction of leaf features has important theoretical and practical significance in biological ecosystems and leaf retrieval. With the development of computer and image processing technology, automatic high-precision extraction of leaf structure has become an emerging interdisciplinary subject. For extracting leaf teeth characteristics, this study followed the biological leaf structure classification system; we focused on describing and extracting leaf teeth by identifying factors such as number of orders of leaf teeth, tooth spacing, number of teeth, sinus shape, and tooth shape. The experimental results showed that the leaf tooth description and measurement method proposed in this study can provide an effective way for a refined description of leaf features and leaf classification; however, the method is only applicable for plants with leaf teeth. For plants without leaf teeth, other features of leaves will need to be considered, which will also be the direction of our work in the next stage. For example, we will explore a method of automatic extraction of leaf-related features through image processing according to the description and definition of leaf organization, glands, petiole, and other features in the Manual of Leaf Architecture. These scientific and canonical leaf characterization data can be used to correlate and contrast plants at different times, thereby, inferring the evolutionary history of plants.

## Author Contributions

Conceived and designed the experiments: HW DT YT. Performed the experiments: DT HW HYZ. Analyzed the data: CL YT. Contributed reagents/materials/analysis tools: CL HYZ. Wrote the paper: HW DT. Provided editorial suggestions and revisions: YT.

